# Metagenomic analysis of a blood stain from the French revolutionary Jean-Paul Marat (1743-1793)

**DOI:** 10.1101/825034

**Authors:** Toni de-Dios, Lucy van Dorp, Philippe Charlier, Sofia Morfopoulou, Esther Lizano, Celine Bon, Corinne Le Bitouzé, Marina Alvarez-Estape, Tomas Marquès-Bonet, François Balloux, Carles Lalueza-Fox

## Abstract

The French revolutionary Jean-Paul Marat (1743-1793) was assassinated in 1793 in his bathtub, where he was trying to find relief from the debilitating skin disease he was suffering from. At the time of his death, Marat was annotating newspapers, which got stained with his blood and were subsequently preserved by his sister. We extracted and sequenced DNA from the blood stain and also from another section of the newspaper, which we used for comparison. Results from the human DNA sequence analyses were compatible with a heterogeneous ancestry of Marat, with his mother being of French origin and his father born in Sardinia. Metagenomic analyses of the non-human reads uncovered the presence of fungal, bacterial and low levels of viral DNA. Relying on the presence/absence of microbial species in the samples, we could cast doubt on several putative infectious agents that have been previously hypothesised as the cause of his condition but for which we detect not a single sequencing read. Conversely, some of the species we detect are uncommon as environmental contaminants and may represent plausible infective agents. Based on all the available evidence, we hypothesize that Marat may have suffered from a fungal infection (seborrheic dermatitis), possibly superinfected with bacterial opportunistic pathogens.

## 1 Introduction

Jean-Paul Marat (1743-1793) was a famous French physician, scientist and journalist, best known for his role as Jacobin leader during the French Revolution. Marat’s parents were Giovanni Mara, born in Cagliari, Sardinia, who later added a “t” to his family name to give it a French feel and Louise Cabrol, a French Huguenot from Castres. Marat was stabbed to death in his bathtub by the Girondist’ supporter Charlotte Corday on July 13^th^, 1793 (Figure 1a). Upon his death, his sister Charlotte Albertine kept two issues of Marat’s newspaper *l’Ami du Peuple* (nº506 and nº678, published on June 30^th^, 1791 and August 13^th^, 1792, respectively), which he was annotating the day of his assassination and that got stained with his blood (Figure 1b). Albertine gave the issues to the collector François-Nicolas Maurin (1765-1848) in 1837. After his death, as explained by a handwritten note by writer Anatole France dated from October 10th, 1864, the two issues ended up in the possession of baron Carl De Vinck who in 1906 donated them to the Département des Estampes, Bibliothèque National de France, in Paris (see notice in the Catalogue Général: https://catalogue.bnf.fr/ark:/12148/cb40261215w).

**Figure 1:**
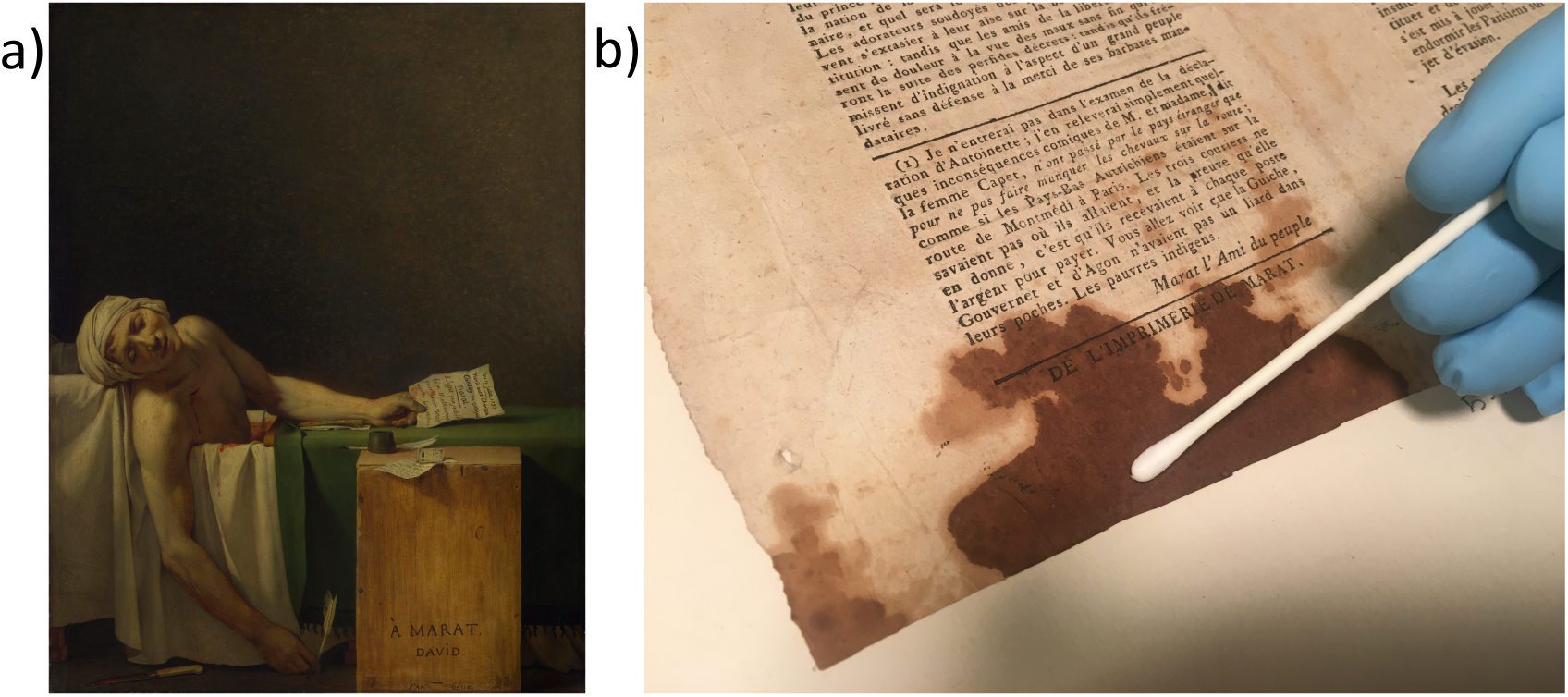
a) “La mort de Marat”; portrait of Jean-Paul Marat after his assassination, by Jacques-Louis David (1793). Preserved at Musées Royaux des Beaux-Arts de Belgique, Brussels. b) Sampling the page of l’*Ami du Peuple* stained with Marat’s blood that has been analysed.

Marat’s health during the last years before his assassination is shrouded in mystery. He suffered from a severe itching skin disease from which he found some relief by spending most of his time in a medicinal bathtub over which he placed a board to use as a writing desk. His condition, which he attributed to his stay in the sewers of Paris while hiding from his political enemies, has been the subject of numerous medical debates and has been alternatively attributed to scabies, syphilis, atopic eczema, seborrheic dermatitis or dermatitis herpetiformis (1–5), the latter as a potential manifestation of celiac disease (6). It has been suggested that his condition affected his character and turned it more violent (1).

With the intention of shedding light on these issues, we retrieved two samples from one of the newspapers stained with Marat’s blood, one sample from the blood stain and a second one from a non-stained area in the upper corner of the paper, to be used as a comparison. A principal concern was to use a non-destructive approach to explore Marat’s genomic footprint; therefore, the samples were taken with forensic swabs. The DNA extracted from both samples was used to build genomic libraries that were subjected to second-generation sequencing using the Illumina platform. DNA reads where subsequently classified, separating the human reads – most likely deriving from Marat’s blood – from those assigned to microbial species. The analysis of both sets of DNA sequences allowed characterisation of Marat’s ancestry as well as identification of the potential pathogens responsible for his debilitating skin condition.

## 2 Material and methods

### 2.1 DNA extraction and sequencing

Forensic swabs were obtained from one of the newspapers Marat was annotating at the time of his assassination (Figure 1). One swab was taken from the blood stain and another from an area of the newspaper without visual evidence of blood. The blood swab was extracted with a buffer composed of 10 mM TrisHCl, 10 mM EDTA, 2 mM SDS, 50 mM DTT; proteinase K was added after one hour incubation. The extract was subsequently concentrated and purified using a Qiagen column kit. DNA extraction from both swabs was performed together with extraction blanks (no sample). A total of 35 ul of each sample was used for library preparation following the BEST protocol (7). Libraries were quantified using BioAnalyzer and sequenced by HiSeq 4000 (Illumina). Library blanks were also performed for each library batch. We generated 568,623,176 DNA reads from the blood stain, of which 74,244,610 reads mapped to the human reference genome (Table S1). From these, we retrieved a complete human mitochondrial (mtDNA) genome at a mean depth of coverage of 18.156X and a nuclear genome at 0.125X.

### 2.2 Mapping and variant calling

Raw sequences adapters were removed using *Cutadapt* (8). Reads were then aligned against the Human Reference genome (GRCh37/hg19) and the mitochondrial reference genome (rCRS) as well as for a set of microbial candidates using *BWA* v.07.3 (9) and Bowtie2 (10). We employed two aligners as mapping sensitivity of different aligners can vary between different samples when working with aDNA (11). Duplicate reads were discarded using *Picard* tools (12). Unique mapped reads were filtered for a mapping quality equal of above 30 (Table S1). All mapped sequences (human nuclear, human mitochondrial and microbial) were assessed for post-mortem damage patterns at the ends of reads using *MapDamage* v.2 (13), which can be used as a sign of historic authenticity over modern contamination (Fig. S1). Post-mortem damage signals were also obtained for each read using *pmdtools* (14) (Fig. S2). Mapping statistics including the depth of coverage were recorded using *Qualimap* (15). Due to the low coverage of the human sample, we performed a pseudo-haploid calling approach, common to the processing of aDNA, using the *SAMtools Pileup* tool (16). This data was then merged with the Human Origins dataset for its use in population genetics analyses (17,18).

### 2.3 Modern DNA Contamination

*Schmutzi* was used to estimate the amount of modern DNA contamination in the mitochondrial (mtDNA) genome (19) likely deriving from the DNA of those who have handled the newspaper in the years following Marat’s death. We identified mitochondrial contamination based on the inferred deamination patterns as 52.5% +/- 4.5% with the full haplogroup profiles provided in Table S2. This allowed the modern DNA sequences to be delineated from the ancient DNA sequences using *Jvarkit* and a custom script (20) by selecting the human reads with mismatches in their first or last three nucleotides. This reduced the amount of modern mtDNA contamination to 0-0.1%. The depth of coverage was recorded using *Qualimap*.

We also independently estimated the amount of contamination based on the heterozygous sites in the X chromosome using *angsd v0.925-21* (21,22). We obtained an estimate of 3.2% modern contamination. As with the mtDNA genome we filtered reads with mismatches in the first or last three nucleotides, taking forward only those reads for additional population genetics analyses. After applying both filters, the resultant mean depth of coverage for the mitochondrial genome was 4.038x and 0.029x for the nuclear genome.

### 2.4 Uniparental Markers and sex Determination analyses

The mtDNA haplogroup was determined using *SAMtools* pileup tool calling the positions defined in the *Phylotree* database (23). We used a genome browser (*IGV*.v2.4.14) to study the genomic context of each possible SNP (24). Only those SNPs that were present in two or more reads, and those which were not located at the ends of the reads, were considered. The contamination was estimated by calculating the ratio of discordant reads at haplogroup-diagnostic positions. Molecular sex was assigned with Ry_compute (22), a script designed for the sex identification of low coverage individuals (Fig. S3).

### 2.5 Population Genetics Analysis

Principal Component Analysis (PCA) was performed using *SmartPCA* in *EIG* v6.0.1 with a subset of modern individuals from the Human Origins dataset (25). This subset contained 434 present-day Europeans and 616,938 autosomal SNPs, plus our sample (Figure 2). The Marat sample was projected using the option *lsqproject*. We also considered the Marat sample projected into an expanded dataset of West Eurasian populations (Fig. S4). As projected individuals’ components tend to 0, we also carried out a control analysis using Han Chinese, French and Marat (Fig. S5). The results were visualised using the R package *GGplot2* (26). This dataset confirmed that Marat is not artefactually placed at the centre of the plot.

**Figure 2:**
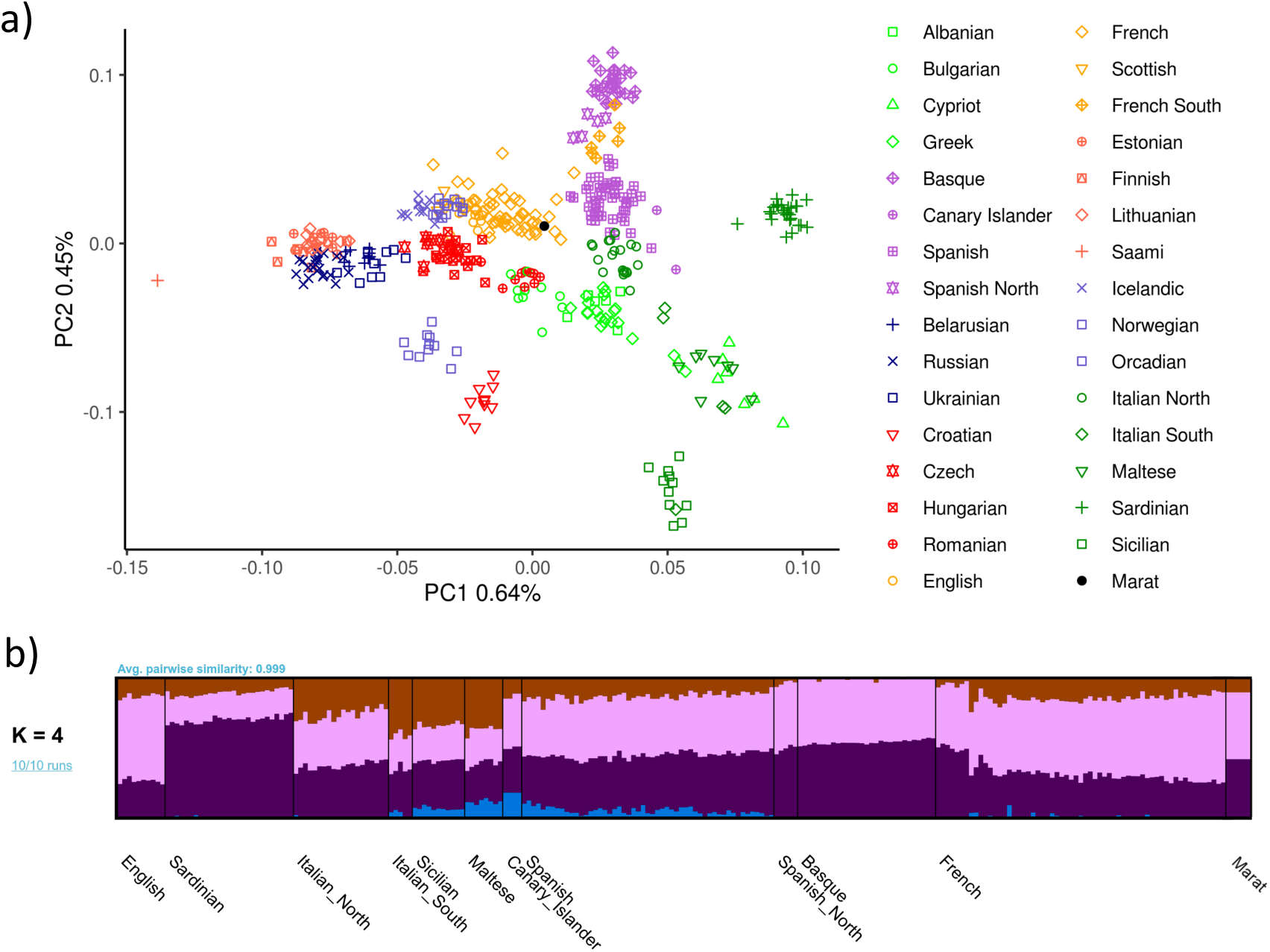
a) Principal Component Analysis (PCA) of modern human European populations with Marat’s ancient DNA reads projected. Symbols provide the country and region, where provided, as given in the legend at right. b) Admixture analysis with modern European samples and Marat. Both analyses are coherent with Marat’s suggested French and Italian combined ancestry.

To formally test the relationship of the Marat sample to relevant geographic regions we calculated *f4* statistics of the form *f4*(Mbuti, Marat; X. Y) where X and Y are tested for combinations of possible ancestral sources: Sardinian, French, English, Italian_North, Basque, Spanish. *f4* values were calculated in qpDstat of AdmixTools v.5.0 (27) with statistical significance assessed through Z-scores following jack-knife resampling (Table S3). This statistic tests the covariance in allele frequency differences between an African out-group (Mbuti) and Marat relative to the clade formed by X and Y. Positive values of *f4* indicate a closer affinity of Marat to Y relative to X, with negative values indicating a closer relationship of Marat to X relative to Y.

We additionally ran an unsupervised clustering analysis using *ADMIXTURE* v1.3 and another subset of the Human Origins dataset (28). This subset included 881 individuals from Europe, West Asia and North Africa typed over 616,938 shared autosomal SNPs. We filtered the dataset by removing SNPs in high linkage disequilibrium using PLINK.v1.9 (29), removing all SNPs with a r^2^ threshold of 0.4 within a 200 SNP sliding window, advancing by 50 SNPs each time. We performed the clustering analysis using K values ranging from 1 to 10, with 10 replicates for each value of K. We selected K according to the lowest cross-validation error value (K=4). The *ADMIXTURE* results at K=4 were visualised using Pong (30) (Figure 2).

### 2.6 Metagenomic analysis

We first removed adapters and merged the paired-end reads into longer single-end sequences using AdapterRemoval v2 (31). We removed PCR duplicates with exact sequence identity using dedupe from the BBMap suite of tools (https://sourceforge.net/projects/bbmap/). We subsequently used the default preprocessing pipeline designed for metaMix which consists of removing human and rRNA sequences using bowtie2 followed by megaBLAST, as well as low quality and low complexity reads using prinseq (32) (-lc_method dust -lc_threshold 7 - min_qual_mean 15). The number of reads filtered at each step are provided in Table S4. We screened the remaining high quality DNA reads for the presence of possible pathogens using both KrakenUniq (33) against the Kraken database compiled in Lassalle et al 2018 (34) and metaMix (35) using megaBLAST and a local custom database consisting of the RefSeq sequences of bacteria, viruses, parasites, fungi and human, as of July 2019. KrakenUniq was run with default parameters. The metaMix-nucleotide mode was run with the default read support parameter of 10 reads was used (Table S5) and the default number of 12 MCMC chains. The number of the MCMC iterations is automatically calculated by metaMix based on the number of species to explore for each dataset, resulting in 10,000 iterations for the blood sample and 3,230 iterations for the paper swab.

The relative proportion of reads assigned to different species by KrakenUniq and metaMix was highly correlated; R^2^=0.94 and R^2^=0.82, for the blood stain and the unstained paper, respectively (Fig. S6). However, metaMix tended to assign a higher number of reads to individual species, closer to the number found by mapping directly to the microbial genomes and we observed important discrepancies for the number of reads assigned to some of the species (Table S6). Additionally, metaMix results for both the blood stain and the unstained paper consisted of fewer species compared to KrakenUniq, even when the same read support threshold was applied to KrakenUniq, indicating increased specificity due to the MCMC exploration of the species space, that comes at an increased computational cost.

In order to compare the accuracy of the two assignment tools, we further explored the presence of clinically relevant species by mapping the quality-filtered subset of reads (Table S2) used in metagenomic assignment against the reference genomes of different candidate genera of fungi and bacteria using *bowtie2* (10) and *BWA* v.07.3 (Fig. S7-S13). For all reads mapping to individual reference genomes, mapDamage v2 (13) was also run to assess evidence of nucleotide mis-incorporation characteristic of post-mortem damage. These mapping results were systematically supporting the metaMix assignments over those obtained with KrakenUniq (Table S6). This led us to rely on metaMix for all metagenomic assignments presented in the paper.

Besides testing for the presence and absence of species, we tested whether some microorganisms were overrepresented in the blood stain compared to the unstained section of the paper using a one-sided binomial test and a significance threshold of 0.95 (Table S5).

As an additional control, we also conducted metagenomic analysis of two publicly available ancient metagenomes obtained from parchment of comparable age to the Marat newspaper (36). We followed the same pre-processing pipeline described for the Marat samples, first removing adapters and PCR duplicates before employing the default metaMix pre-processing pipeline, this time removing reads that mapped to either the human or sheep, cow and goat reference genomes. As before, metaMix-nucleotide mode was run with with a read support parameter of 10 reads and with 12 MCMC chains x 2,325 and 6,130 iterations respectively for ERR466100 and ERR466101. We provide the breakdown of read filtering steps in Table S7 and our raw metaMix results in Table S8.

### 2.7 Phylogenetic analysis

In the case of *Malassezia*, a phylogenetic analysis of the mitochondrial DNA genome with available modern strains on the Short Read Archive (SRA) was performed. We called variant positions using *GATK UnifiedGenotyper* (37) and generated a Maximum Likelihood tree using *RAxML-NG* specifying a GTR substitution model and 100 bootstrap resamples (38). The tree was rooted with *M. globosa* (Fig. S10).

We also conducted a phylogenetic analysis for *C. acnes*, combining our historical strain with all *C. acnes* genomes deposited in the SRA covering the reference at an average depth >10x, and with *C. namnetense* as an outgroup (SRR9222443). The only *C. acnes* genomes sequenced at medium to high depth are those reported by Gomes et al 2017 (39). A Maximum Likelihood tree was generated over the 21,751SNP alignment using RAxML-NG (Fig. S13) and clonal complexes and phylotypes were assigned based on the PubMLST *C. acnes* definitions database (https://pubmlst.org/bigsdb?db=pubmlst_pacnes_seqdef).

## 3 Results

### 3.1 Human ancestry analysis

We generated 568,623,176 DNA reads from the blood stain, of which 74,244,610 reads mapped to the human reference genome (Table S1). From these, we retrieved a complete human mitochondrial (mtDNA) genome at a mean depth of coverage of 4.038x and the nuclear genome at 0.029x (Table S1). The predominant mtDNA haplotype was H2a2a1f, although we found evidence of some additional mtDNA sequences, notably a K1a15 haplotype. The ratio of sexual chromosome to autosomal DNA reads indicated that the sample donor was male (Fig. S3).

The human DNA reads showed evidence of post-mortem deamination occurring in 1% of the ends of sequencing reads, indicating authentic ancient DNA damage (Fig. S1-S2). This is similar to the degree of damage that has been observed in aDNA obtained from other human specimens of a similar age (40). For further analyses we selected only those reads that displayed C to T or G to A substitutions at the 5’ or 3’ end, respectively. After this procedure, the degree of mitochondrial contamination was reduced to 0-0.01%.

To explore the ancestry of Marat in the context of modern European populations, we performed Principal Component Analysis (PCA) (Figure 2a and Fig. S4-5) and unsupervised clustering in ADMIXTURE (Figure 2b). Our sample projected among modern French individuals sampled from France in the population genetic analyses. This result is broadly compatible with proposed hypotheses relating to the ancestry of Marat (2). *f4* statistics suggest a closer affinity of Marat to modern Italian, English, Sardinian, Basque and French populations relative to those from Spain (Table S3). However, these trends are subtle and we note that mixed ancestries are difficult to discern, especially when only limited genetic data is available.

### 3.2 Metagenomic analysis

We conducted metagenomic species assignments with the 9,788,947 deduplicated, quality controlled and low complexity filtered DNA reads (combined merged and non-merged) that did not map to the human genome (see Methods and Table S4). We used metaMix (35), a Bayesian mixture model framework developed to resolve complex metagenomic mixtures, which classified ∼9% of the non-human reads into 1,328 microbial species (Table S5). The species assignments were replicated with KrakenUniq (33), which led to largely consistent, if less accurate, results (∼7% classified into 3,213 species, Fig. S6, Table S6). Thus, we relied on the metaMix species assignments throughout the paper, unless stated otherwise.

We detected the presence of a wide range of microorganisms, including some expected to develop on decaying cellulose and/or dried blood, but also others recognized as opportunistic human pathogens from the following bacterial genera: *Acidovorax, Acinetobacter, Burkholderia, Chryseobacterium, Corynebacterium, Cutibacterium, Micrococcus, Moraxella, Paraburkholderia, Paracoccus, Pseudomonas, Rothia, Staphylococcus, Streptococcus* and the fungal genera *Aspergillus, Penicillium, Talaromyces* and *Malassezia* as well as HPV (type 179 and type 5) and HHV6B viruses, albeit the latter supported by a very low numbers of reads (Table S5-S6). Some of the DNA reads, notably from *Aspergillus glaucus, Cutibacterium acnes, Malassezia restricta* and *Staphylococcus epidermis* showed typical misincorporation patterns that are considered indicative of these sequences being authentically old (Fig. S7).

We additionally sequenced the swab taken from the unstained paper sample. In this case, only 96,252 pairs of reads were obtained (56,616 merged, 25,712 non-merged, 35,216 deduplicated and filtered combined merged and non-merged), with 52% of the reads that could be classified with metaMix into 66 species and 36% with KrakenUniq into 374 species, respectively (see Methods and Table S4). Although very little DNA could be retrieved from the section of the document that had not been blood-stained, we tried to identify microorganisms that were statistically significantly over-represented in the blood stain relative to the unstained paper. Amongst these and besides, as expected, *Homo sapiens*, different species of *Aspergillus* and *Acinetobacter* were significantly overrepresented in the blood stain (Table S5). It remains questionable however whether the unstained paper represents a suitable negative control given that the newspaper had been extensively manipulated by Marat. Significant over-representation of *Aspergillus spp.* and *Acinetobacter spp.* in the blood stain relative to the rest of the document could also be due to the blood providing better conditions for the growth of iron-limited microbes. Indeed, *Aspergillus spp.* and *Acinetobacter spp.* are commonly found in the environment but are also grown in blood agar. As such, it is plausible that these represent *post-mortem* contaminants. Indeed, for *Acinetobacter spp.* we identified no post-mortem damage pattern.

Metagenomic analysis of historical samples can be challenging as the resulting microbial communities typically comprise an unknown mixture of endogenous species as well as contaminants, both contemporary and modern. To mitigate this problem, we relied on a ‘differential diagnostics’ approach (Table 1), where we specifically tested for the presence of reads from pathogens that could plausibly have led to Marat’s symptoms, most of which have been previously hypothesised in the literature (1–5). Such a differential approach is more stringent than the standard approach in clinical diagnostics aiming to identify the full list of microbes present in the samples after enforcement of a read-number threshold (41,42). Our approach allows limiting the number of species to be tested to a small list of plausible candidates. Second, the lack of detection of even one read from a focal microbial species by direct mapping falsifies the null hypothesis that it was not involved in the disease.

**Table 1:** List of diseases tested for associated agents and presence in the blood stain and the unstained paper samples. The following symbols denote the abundance of reads for each infectious agent tested. ✓: present; ✓✓: top ten; ✓✓✓: top hit; ✘: absent.

| Disease | Pathogen | Blood | Unstained paper |
| --- | --- | --- | --- |
| Syphilis | *Treponema pallidum* | ✕ | ✕ |
| Scrofula (tuberculosis) | *Mycobacterium tuberculosis*<sup>1</sup> | ✕ | ✕ |
| Leprosy | *Mycobacterium leprae* | ✕ | ✕ |
| Diabetic candidiasis (thrush) | *Candida* sp. | ✕ | ✕ |
| Scabies | *Sarcoptes scabiei* | ✕ | ✕ |
| Seborrheic dermatitis | *Malassezia* sp. | ✓✓ | ✓ |
| Atopic dermatitis | *Staphylococcus aureus* | ✓ | ✕ |
| Severe acneiform eruptions | *Cutibacterium acnes* | ✓✓✓ | ✓✓ |

<sup>1</sup> Scrofula can also be caused by other Mycobacteria in particular *M. scrofulaceum* and *M. avium intracellulare*, which are also absent from both samples.

We did not identify a single sequencing read in either the blood stain or the unstained paper for the agents of syphilis, leprosy, scrofula (tuberculosis) and diabetic candidiasis (thrush) (Table 1, Table S5). We additionally tested for scabies, which is caused by burrowing of the mite *Sarcoptes scabiei* under the skin. Since the metagenomic reference database did not include arthropod genomes, this was tested separately by blasting all the non-human reads against the *Sarcoptes scabiei* genome (GCA_000828355.1). Again, we detected not a single read matching to *Sarcoptes scabiei*, which makes scabies an implausible cause for Marat’s skin disease (Table 1, Table S5).

Conversely, metaMix recovered 15,926 and 83 filtered DNA reads from the blood stain and the unstained paper respectively, assigned to *Malassezia restricta* a fungal pathogen causing seborrheic dermatitis, which has been previously hypothesized as one of the most plausible causes for Marat’s condition (1–4). Direct mapping of all reads to *M. restricta* (GCA_003290485.1) resulted in 19,194 reads from the blood stain dataset mapping over 17.17% of the reference genome. KrakenUniq failed to identify *M. restricta*, instead assigning 627 reads sequenced from the blood stain to *M. sympodialis*. However, further analysis of the *Malassezia* reads based on genome mapping pointed to most (80.3%) being uniquely assigned to *M. restricta* rather than *M. sympodialis* (Fig. S8). This allowed us to reconstruct a complete *M. restricta* mtDNA genome at 0.84X coverage. The *Malassezia* reads were evenly distributed along the full genome assembly supporting no mixing or misclassification of the species (Fig. S9).

We placed our Marat *M. restricta* mitochondrial genome in phylogenetic context by building a maximum likelihood phylogeny including our historical strain and available present-day mtDNA *M. restricta* genomes. Although the total number of samples is small, the fact that the *M. restricta* mtDNA molecule recovered from Marat’s blood is placed basal to modern strains (Fig. S10) and exhibits some post-mortem damage (Fig. S5) further support its authenticity.

We also recovered 587 filtered reads assigned by metaMix to *Staphylococcus aureus* in the blood stain but none in the reads obtained from the unstained paper. The differential representation in the two samples is not significantly different due to the far lower number of reads in the unstained sample (Table S5). Although a common commensal, *S. aureus* is also a frequent human pathogen and the leading cause of atopic eczema. In order to confirm the metagenomic assignments to *S. aureus*, we mapped the raw microbial reads to a series of reference genomes from various species in the *Staphylococcus* genus. This allowed us to identify 888 reads mapping against the *S. aureus* reference genome, out of which 758 uniquely mapped to *S. aureus* (Fig. S11). The presence of *S. aureus*, but with a relatively low number of reads, may be compatible with a secondary infection by *S. aureus* rather than *S. aureus* being the initial cause of Marat’s condition. Alternatively, Marat, or someone who also handled the newspaper, could have carried *S. aureus* as a skin commensal.

The most prevalent microbial species in the blood stain was *Cutibacterium acnes* (formerly *Propionibacterium acnes* (43)), which was also present in the unstained paper (Table S5). *C. acnes* is largely a commensal and part of the normal skin biota present on most healthy adult humans’ skin, including in association with *S. epidermis* which we also observe in our sample (Table S5-S6, Fig. S11) (44). *C. acnes* is also a frequent contaminant in metagenomic samples (45,46). However, *C. acnes* can also be involved in severe acneiform eruptions (47) and we cannot exclude the possibility that it could have contributed to Marat’s condition. 86,019 reads mapped to the *C. acnes* reference genome (GCF_000008345.1), yielding an alignment of 3.4X average coverage (Fig. S12) and exhibiting modest post-mortem damage (Fig. S7).

A phylogeny of Marat *C. acnes* with a collection of publicly available modern strains (39,46) places our historic genome on a short branch falling basal to Type I strains, supporting its age and authenticity (Fig. S13). This phylogenetic placement suggests our Marat strain falls into *C. acnes* phylotype I (*C. acnes subsp. acnes*) rather than II (*C. acnes subsp. defendens*). Whilst our Marat strain does not cluster with phylotype Ia, the type more commonly associated with skin surface associated acne vulgaris (48), its position, basal to Type Ib strains cannot exclude its involvement in soft or deep tissue infections (49).

Delineating contaminants and commensals from plausible pathogens remains challenging from this type of data source, in particular due to the absence of a suitable control. To alleviate this issue, we conducted full taxonomic assignments of two ancient metagenomes generated from historical parchment samples dating to the 17^th^ and 18^th^ centuries (PA1 and PA2 respectively) (36). Although these samples were obtained from livestock (ruminant) skins whereas we are working with cellulose paper, we anticipate that they may have been used and handled in a comparable way to the newspaper Marat was annotating. In this way they represent what can be considered as the most biologically comparable ancient metagenomes available to date. An equivalent metaMix analysis applied to these filtered sequencing reads (Table S7) identified not a single read assigned to *M. restrica, S. aureus* or *C. acnes* (Table S8). We therefore do not systematically expect a significant number of reads for the three species we suggest as most plausible candidates for Marat’s condition.

## 4 Discussion

Over the last decade, ancient-pathogen genomics has made great progress by borrowing technological advances originally developed for the study of human ancient DNA (50,51). Although most microbial data has been secondarily generated from the sequencing of ancient human bones or teeth (51–54) other, rare samples, such as preserved tissues (55,56) or microscope slides from antique medical collections have been analysed (50,57). We are aware of no previous attempt to leverage ancient DNA technology to diagnose infections in historical characters, despite previous sequencing of remains from other prominent historical figures such as King Richard III and the putative blood of Louis XVI (58,59).

In this work we analysed both human and ‘off-target’ microbial reads to shed light on an important historical figure of the French Revolution and his skin condition. Due to the loss of Marat’s remains after their removal from the Panthéon in February 1795, the paper stained with his blood likely represents the only available biological material to study both his ancestry and the cause of his skin condition. Although second-generation sequencing techniques have been applied to the analysis of ancient parchments (60) our work represents the first instance where this methodological approach has been applied to old cellulose paper.

The presence and relative abundance of different microorganisms in the documents Marat was annotating is affected by their endogenous presence as well as contemporary and modern contamination both for the blood and unstained sample. Some microorganisms present in the samples might reflect skin microbiome signatures. Whilst some other microorganisms represent environmental contaminants and are likely unrelated to Marat’s condition. In order to identify the most likely candidates for Marat’s condition we tested a set of proposed diagnoses, which we considered as plausible if we detected at least one read assigned to the causative infectious agent (Table 1).

Potential conditions for which we detected not a single supportive read included syphilis, tuberculosis (scrofula), leprosy, diabetic candidiasis or scabies. We appreciate that absence of evidence for an infectious agent does not constitute incontrovertible evidence of its absence. Moreover, it is not uncommon for metagenomic diagnostics applied to clinical samples to fail to identify reads from the likely infectious agent above the predefined diagnostic threshold, or even fail to detect any read at all (42,43). As such, the absence of reads from a putative pathogen makes it less plausible as the agent of Marat’s condition but does not definitely rule them out.

Conversely, we detected and validated microbial reads for two of the conditions we tested, seborrheic dermatitis (*Malassezia spp.)*, atopic dermatitis (*Staphylococcus aureus*) and cannot exclude severe acneiform eruptions (*Cutibacterium acnes*) as a third, given the age and phylogenetic position of the *C. acnes* genome we obtained. For all three cases, the number of reads would have exceeded the threshold suggested for detection in clinical metagenomic diagnostic (42,43)(41), even when considering the swab from the unstained paper as a control.

The presence of *Malassezia restricta* is of particular interest because this fungus is specialized to live on the skin (61). Although also a common commensal and contaminant in metagenomic studies, *Malassezia* has been described in various skin conditions, including dandruff, atopic eczema, folliculitis and seborrheic dermatitis (62,63). Interestingly, the latter symptoms would fit those described in Marat (5). The *M. restricta* reads we identified were not statistically significantly overrepresented in the blood’s stain relative to the unstained paper, although they could be expected to be present in both samples if someone heavily infected was holding the newspaper. Although we cannot confidently claim the reads in Marat’s blood are directly associated with Marat himself, we do identify post-mortem damage in these reads and a phylogenetic placement in a modern mitochondrial DNA phylogeny consistent with these reads being indeed old (Fig. S7, Fig. S10). We also do not systematically expect the presence of *M. restricta* on parchment of a similar age (Table S8).

Also of possible interest is the widespread presence of *Cutibacterium acnes subsp. acnes*, which although a common commensal or contaminant can also be implicated in severe acneiform eruptions, which constitutes the top hit in the blood sample and falls basal to phylotype I strains currently in circulation. As with *M. restricta*, we do not observe a single *C. acnes* read in two biologically equivalent historic parchment metagenomes (Table S8). *Staphylococcus aureus*, which is frequently detected in cases of atopic dermatitis, is also present in reads obtained from the blood’s stain, although in fairly low number.

Whilst our results do not allow us to reach a definite diagnosis of Marat’s condition, they allowed us to cast doubt on several previous hypotheses and provide, using all the available evidence, some plausible aetiologies. We suggest that Marat could have been suffering from an advanced fungal or polymicrobial infection, either primary or secondary to another condition. Future metagenomic analysis of additional documents in Marat’s possession during his assassination could help confirm the microbial composition found in this study and strengthen these observations.

Our work further illustrates the potential of sequencing technologies for the generation of (meta-)genomic information from difficult, singular samples and opens new avenues to address medical hypotheses of major historical interest.

## Acknowledgments and Funding

This work was supported by Obra Social ‘‘La Caixa’’ and Secretaria d’Universitats i Recerca (GRC2017-SGR880) (T.M.-B. and C.L.-F.), BFU2017-86471-P and PGC2018-101927-B-I00 (MINECO/FEDER, UE) (T.M.-B) and PGC2018-095931-B-100 (MINECO/FEDER, UE) (C.L.-F.). T.M-B. is also supported by a U01 MH106874 grant and Howard Hughes International Early Career and CERCA Programme del Departament d’Economia i Coneixement de la Generalitat de Catalunya. S.M is funded by a Welcome Trust post-doctoral fellowship (206478/Z/17/Z). L.v.D and F.B. acknowledge financial support from the Newton Fund UK-China NSFC initiative (grant MR/P007597/1) and the BBSRC (equipment grant BB/R01356X/1). Full metagenomic reads are available at NCBI under BioProject ID XXXXX.

## Author Contributions

P.C., L.v.D., F.B. and C.L.-F. conceived and designed the study; P.C. and C.L.B sourced the newspaper; C.B., M.A-E. and E.L. developed and performed laboratory analysis; T.d.-D., L.v.D., S.M. analysed data and performed computational analyses; T.d.-D., L.v.D., S.M., F.B., and C.L.-F. wrote the paper with inputs from all co-authors.

