## Supplementary material for "Metagenomic analysis of a blood stain from the French revolutionary Jean-Paul Marat (1743-1793)"

### Supplementary Tables

**Supplementary Table S1:** Mapping metrics of human DNA reads obtained from Marat's blood stain against the human mitochondrial genome (rCRS) and whole human genome (hg19). Unmapped reads following deduplication were taken forward for quality control and filtering (**Table S2**).

| Mapping Statistics | Sequenced reads | Mapped reads | RM duplicates | Quality 30 reads | % duplication | Reads with damage | Coverage Final | Mapped Bases |
|---|---|---|---|---|---|---|---|---|
| **mtDNA** | 568,623,176 | 57,654 | 3,485 | 3472 | 93.96 | 915 | 4.04X | 66,911 |
| **Nuclear** | 568,623,176 | 74,244,610 | 4,686,746 | 4,461,919 | 93.69 | 1,245,497 | 0.03X | 90,301,707 |

**Supplementary Table S2 (external Excel document):** Human mtDNA haplogroup assignments. Alleles and depth are characterised for describing haplogroups positions. Positions with more than two folds of allele depth are highlighted and considered as likely to be part of a defined haplogroup. The majority haplogroup is estimated using the number of likely positions. Contamination estimates are both calculated using schmutzi and from discordant positions in the haplogroups.

**Table S3:** f4 statistics of the form *f4*(Mbuti, Marat; X. Y) where X and Y are tested for combinations of possible ancestral sources: Sardinian, French, Basque, French, Italian_North, Basque. Positive *f4* values indicate a relatively higher affinity of Marat to Y relative to X. Negative *f4* values indicate a relatively higher affinity of Marat to X relative to Y. Z-scores following block jack-knife resampling are provided with a value >|2| considered statistically significant (highlighted in red).

| Combination | | | | *f4* | Z Score |
|---|---|---|---|---|---|
| Mbuti | Marat | Sardinian | French | -0.000017 | -0.061 |
| Mbuti | Marat | French | Sardinian | 0.000017 | 0.061 |
| Mbuti | Marat | English | Basque | -0.000049 | -0.127 |
| Mbuti | Marat | Basque | English | 0.000049 | 0.127 |
| Mbuti | Marat | French | Italian_North | -0.000043 | -0.159 |
| Mbuti | Marat | Italian_North | French | 0.000043 | 0.159 |
| Mbuti | Marat | Sardinian | Italian_North | -0.000061 | -0.189 |
| Mbuti | Marat | Italian_North | Sardinian | 0.000061 | 0.189 |
| Mbuti | Marat | English | Sardinian | -0.000297 | -0.754 |
| Mbuti | Marat | Sardinian | English | 0.000297 | 0.754 |
| Mbuti | Marat | Basque | Sardinian | -0.000247 | -0.799 |
| Mbuti | Marat | Sardinian | Basque | 0.000247 | 0.799 |
| Mbuti | Marat | English | Italian_North | -0.000357 | -0.928 |
| Mbuti | Marat | Italian_North | English | 0.000357 | 0.928 |
| Mbuti | Marat | Basque | Italian_North | -0.000308 | -0.969 |
| Mbuti | Marat | Italian_North | Basque | 0.000308 | 0.969 |
| Mbuti | Marat | English | French | -0.000314 | -0.972 |
| Mbuti | Marat | French | English | 0.000314 | 0.972 |
| Mbuti | Marat | Basque | French | -0.000265 | -1.077 |
| Mbuti | Marat | French | Basque | 0.000265 | 1.077 |
| Mbuti | Marat | Italian_North | Spanish | -0.000713 | -2.697 |
| Mbuti | Marat | Spanish | Italian_North | 0.000713 | 2.697 |
| Mbuti | Marat | Sardinian | Spanish | -0.000774 | -2.888 |
| Mbuti | Marat | Spanish | Sardinian | 0.000774 | 2.888 |
| Mbuti | Marat | English | Spanish | -0.001071 | -3.094 |
| Mbuti | Marat | Spanish | English | 0.001071 | 3.094 |
| Mbuti | Marat | Basque | Spanish | -0.001021 | -3.986 |
| Mbuti | Marat | Spanish | Basque | 0.001021 | 3.986 |
| Mbuti | Marat | French | Spanish | -0.000757 | -4.019 |
| Mbuti | Marat | Spanish | French | 0.000757 | 4.019 |

**Supplementary Table S4:** Number of DNA reads from the blood-stained sample and unstained sample at each filtering steps prior to taxonomic classification. The final read counts for KrakenUniq and metaMix are provided in blue and red respectively.

| **Blood stained sample** | | | |
|---|---|---|---|
| **Filtering step** | **Merged into longer single end reads** | **Non-merged** | **Combined (merged and non-merged)** |
| Deduplicated (bbdedupe) | 11,336,155 | 2,240,430 | 13,576,585 |
| After human removal (bowtie2) | 8,617,745 | 1,505,994 | 10,123,739 |
| After rRNA removal (bowtie2) | 8,578,821 | 1,485,366 | 10,064,187 |
| After QC (prinseq) | 8,486,659 | 1,444,856 | 9,931,515 |
| After human removal (megaBLAST) | 8,370,312 | 1,437,288 | 9,807,600 |
| After rRNA removal (megaBLAST) | 8,356,389 | 1,432,558 | 9,788,947 |
| Reads mapping to nucl DB (megaBLAST) | 1,012,140 | 252,247 | 1,264,387 |
| **Unstained sample** | | | |
| **Filtering step** | **Merged into longer single end reads** | **Non-merged** | **Combined (merged and non-merged)** |
| Deduplicated (bbdedupe) | 18,539 | 111,432 | 129,971 |
| After human removal (bowtie2) | 17,593 | 18,550 | 36,143 |
| After rRNA removal (bowtie2) | 17,488 | 18,550 | 36,038 |
| After QC (prinseq) | 17,352 | 18,454 | 35,806 |
| After human removal (megaBLAST) | 17,276 | 18,410 | 35,686 |
| After rRNA removal (megaBLAST) | 17,156 | 18,060 | 35,216 |
| Reads mapping to nucl DB (megaBLAST) | 8,476 | 9,308 | 17,784 |

**Supplementary Table S5 (external Excel document):** Read classification and species assignment by metaMix of reads obtained from the swab of the unstained paper and blood-stained paper. *P*-values provide those species significantly over-represented in the bloodstained paper compared to the control.

**Supplementary Table S6 (external Excel document):** The number of quality filtered reads mapping to candidate species using a Bowtie2 and BWA pipeline. The number of reads classified by metaMix and KrakenUniq are also provided.

**Supplementary Table S7:** Number of DNA reads from the two historic parchment samples at each filtering steps prior to taxonomic classification. The final read counts for metaMix are provided in red.

| Historic Parchment Samples | | |
|---|---|---|
| **Filtering step** | **ERR466100 (PA1)** | **ERR466101 (PA2)** |
| Raw data | 17006629 | 31493502 |
| After human removal (bowtie2) | 16921807 | 31410793 |
| After rRNA removal (bowtie2) | 16916091 | 31403665 |
| After ruminant removal (bowtie2) | 6232580 | 22161979 |
| After QC (prinseq) | 6157782 | 21951980 |
| After human removal (megaBLAST) | 6093733 | 21753485 |
| After rRNA removal (megaBLAST) | 5056204 | 18517414 |
| After ruminant removal (megaBLAST) | 1557368 | 6618990 |
| Reads mapping to nucl DB (megaBLAST) | 9367 | 43597 |

**Supplementary Table S8 (external Excel document):** Read classification and species assignments by metaMix of reads obtained from two historic parchment samples published in *Teasdale et al.* 2015.

### Supplementary Figures

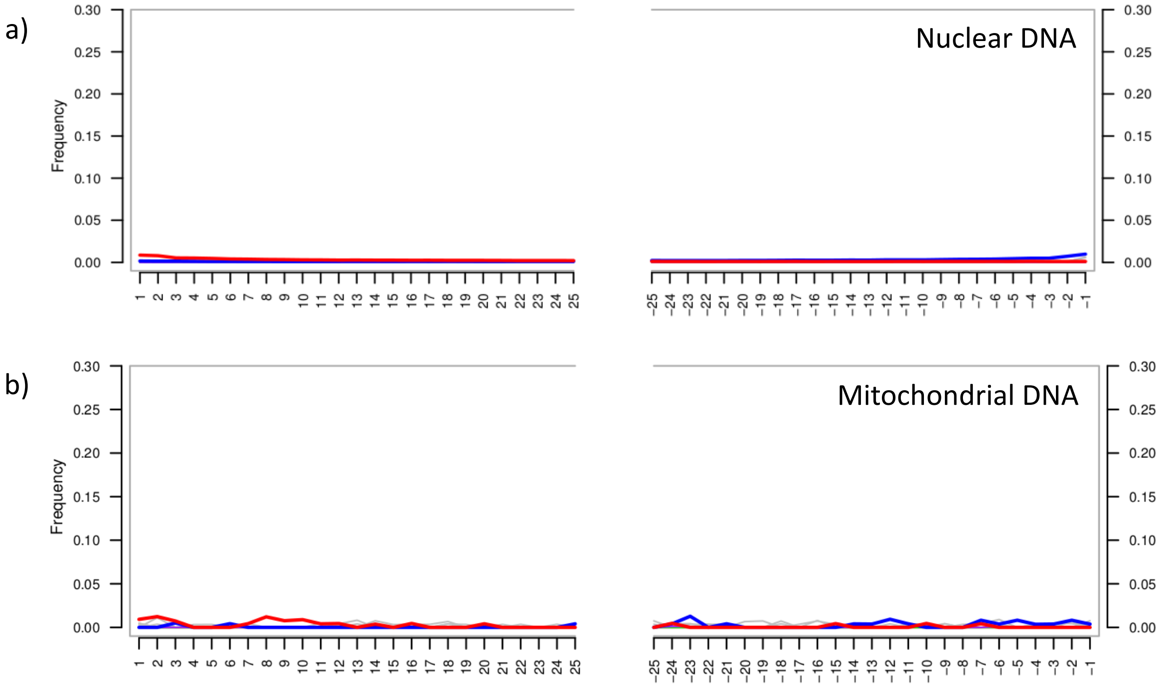


**Supplementary Figure S1: a)** Nucleotide misincorporation patterns at the ends of the human reads obtained from the blood stain sample (0.98% G>A at 3' and 0.86% C>T at 5'). b) Nucleotide misincorporation patterns for human mitochondrial reads only (0.38% G>A at 3' and 0.92% C>T at 5'). In both cases the red line provides the C to T substitution frequency and the blue line provides the G to A substitution frequency from 5' (left) to 3' (right). While the deamination detected is small, we observed that the post-mortem score for each read closely follows that found in a similarly old sample, a 100 year old aboriginal Australian hair sample.

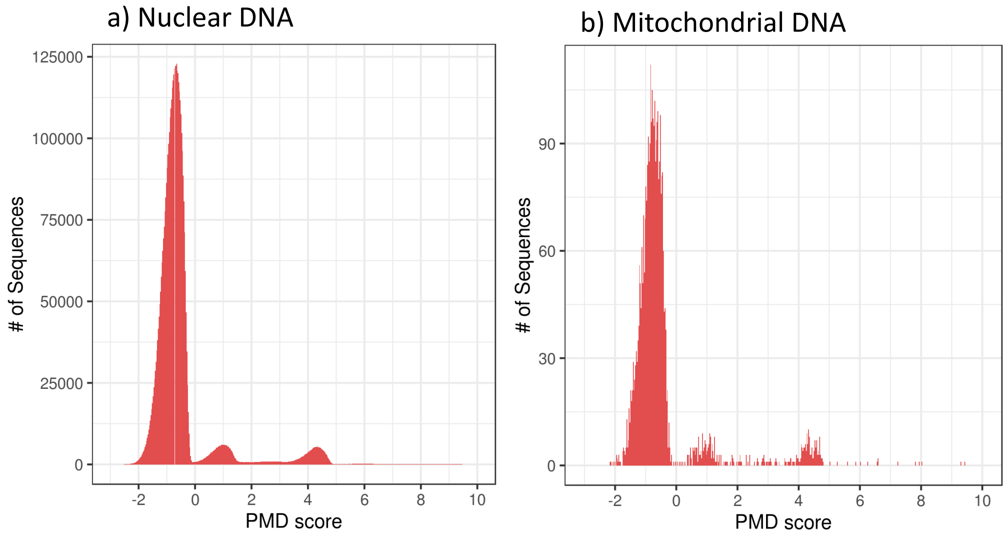


**Supplementary Figure S2:** Post-mortem damage score distributions for the human sequencing reads obtained from the blood-stained paper. a) provides the distribution of scores for the nuclear DNA, b) provides the distribution for the mitochondrial DNA.

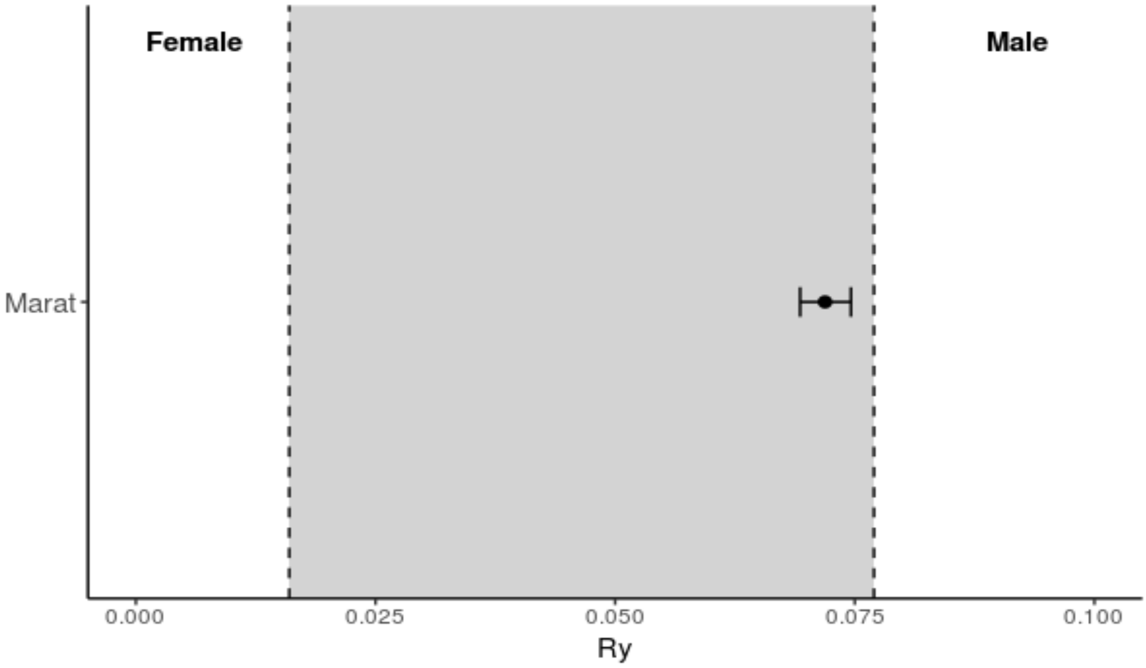


**Supplementary Figure S3:** Sex determination of the human reads inferred from the ratio between the number of reads aligned to the Y chromosome and the total number of reads aligned to both sex chromosomes (Ry). Error bars provide the 95% confidence value. The grey area represents the area where the sex cannot be determined. The sample is incompatible with being a female.

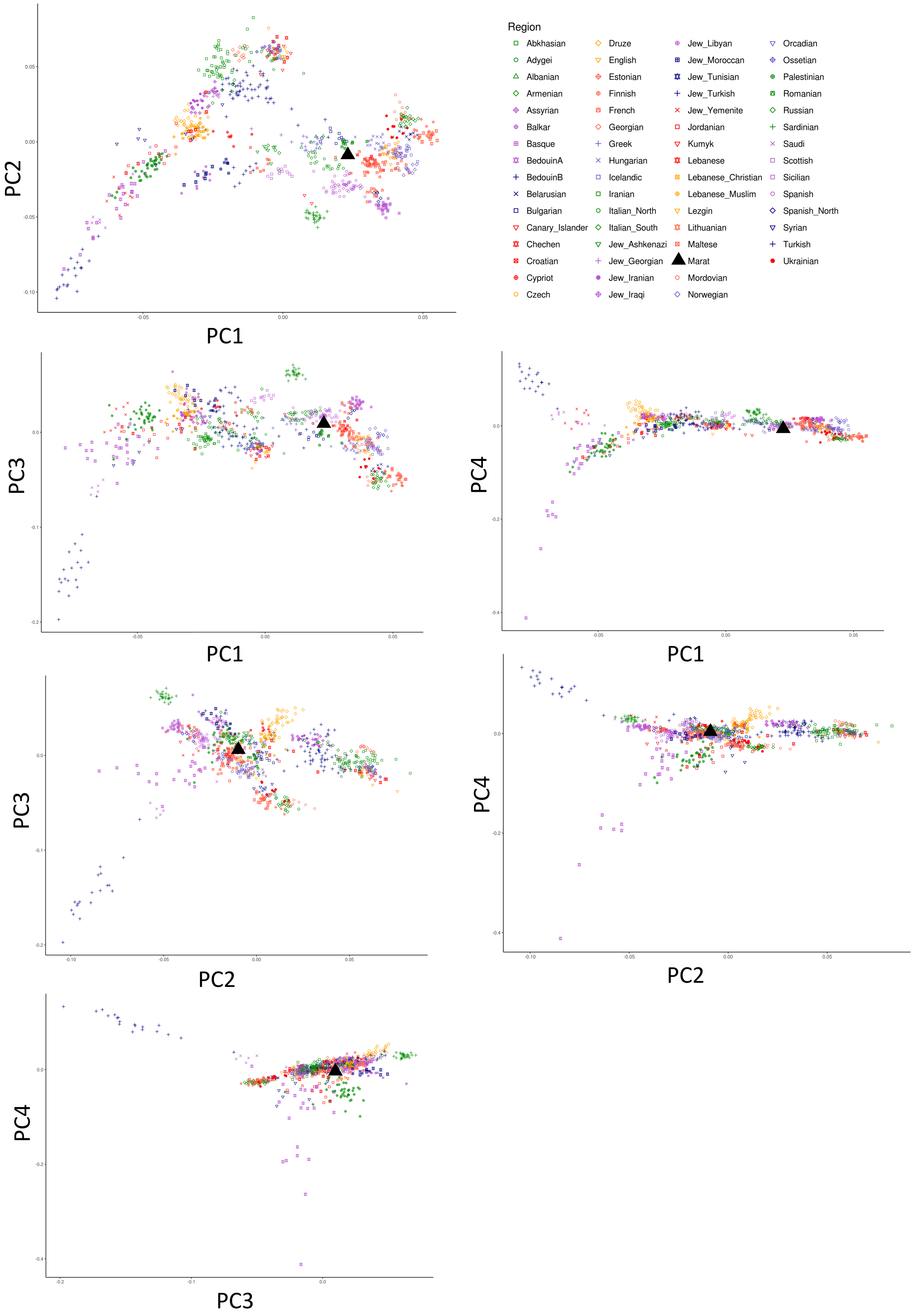


**Supplementary Figure S4:** Principal components plots provided for PC1-4 for West Eurasian populations coloured as per the legend (top-right). The Marat sample (triangle) is projected into PC space in each case using *lsqproject* from *SmartPCA* in *EIG v6.0.1*.

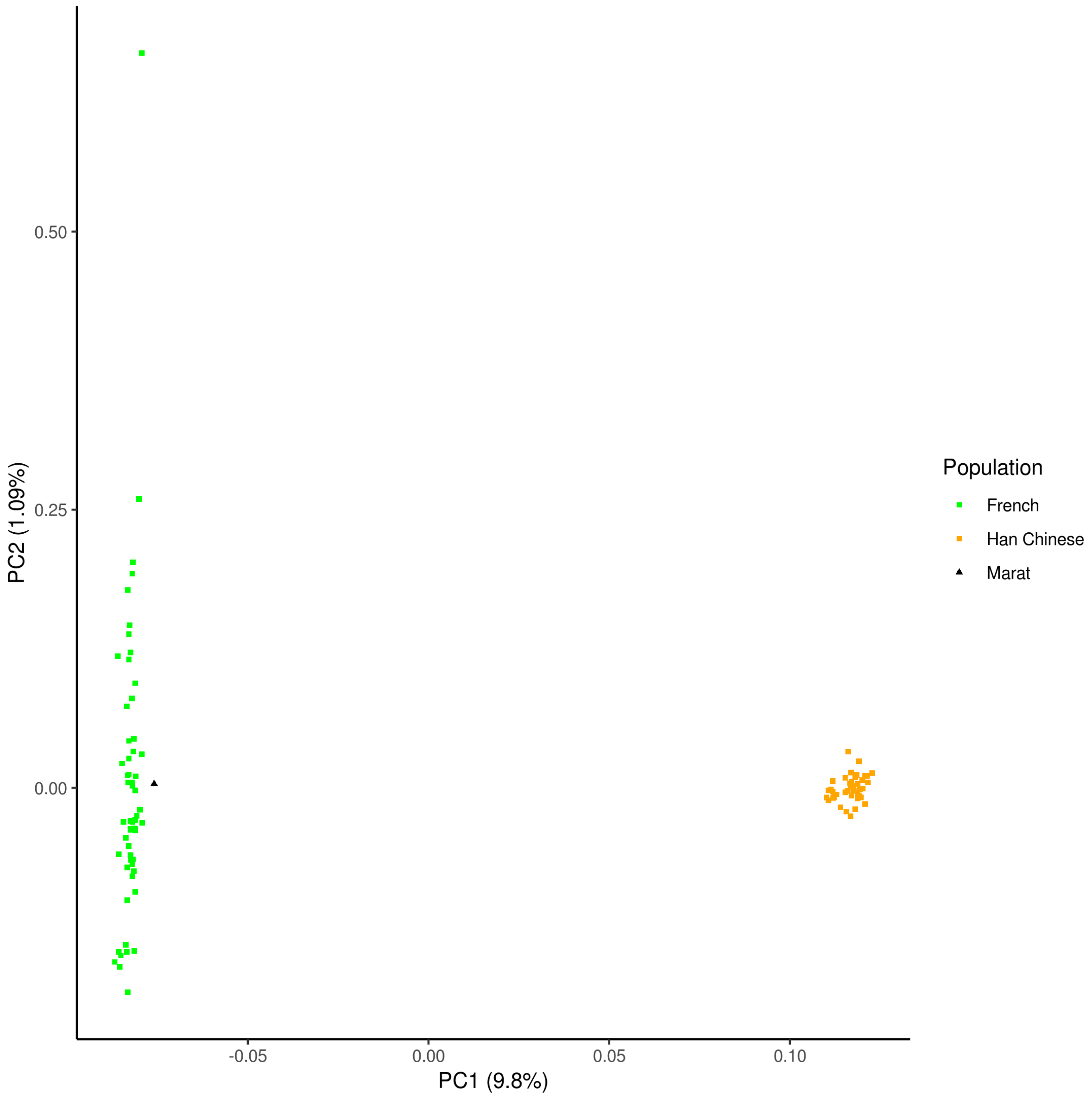


**Supplementary Figure S5:** Principal Component Analysis (PCA) of Marat's sample together with current French and Han Chinese individuals. Marat (triangle) is located at the lower extreme of PC1, clustering with the modern-day French.

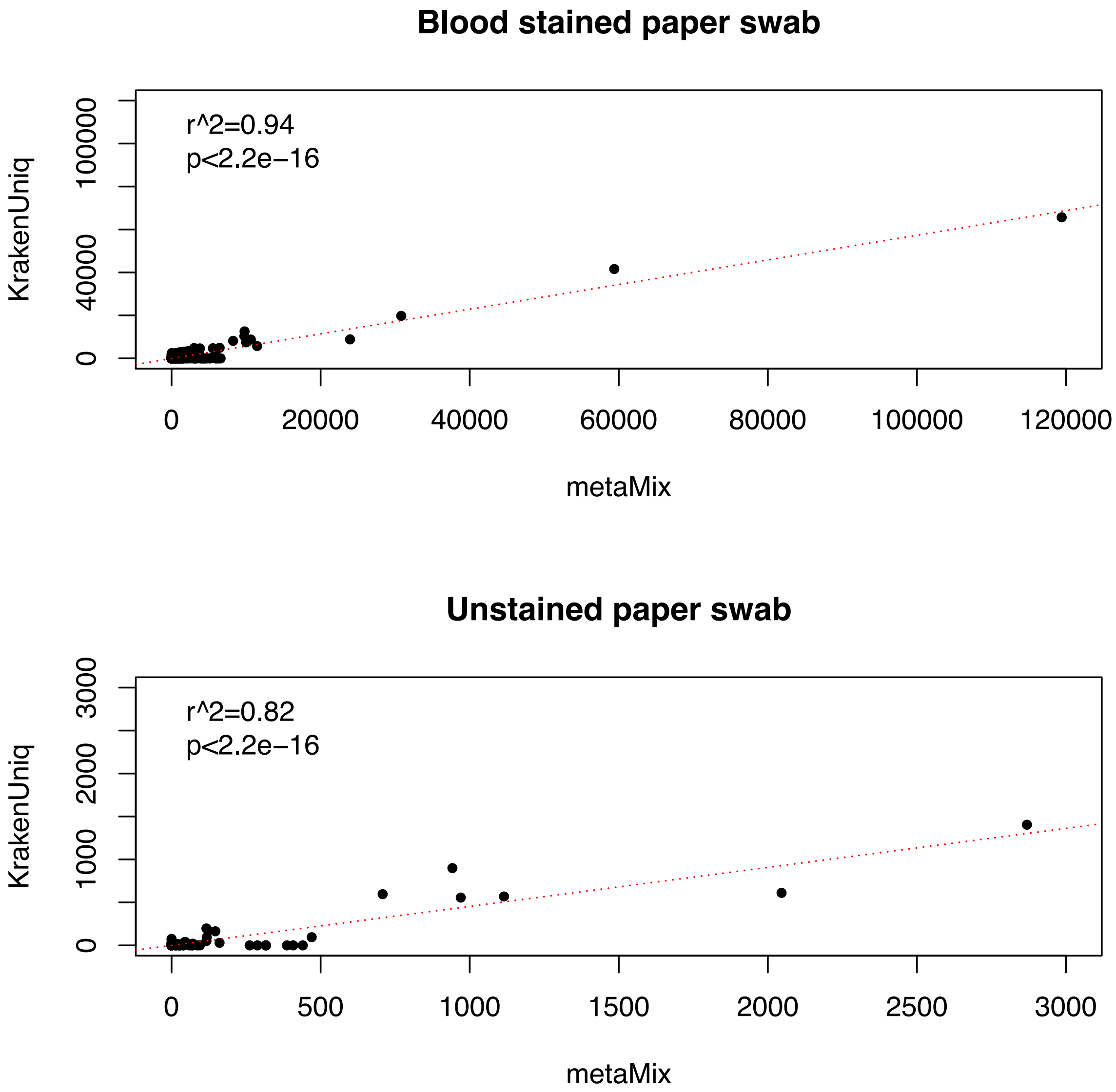


**Supplementary Figure S6:** Linear correlations between the number of reads assigned to individual species by the metaMix (x-axis) and KrakenUniq (y-axis) metagenomic assignment tools for the blood stain (upper panel) and unstained paper (lower panel).

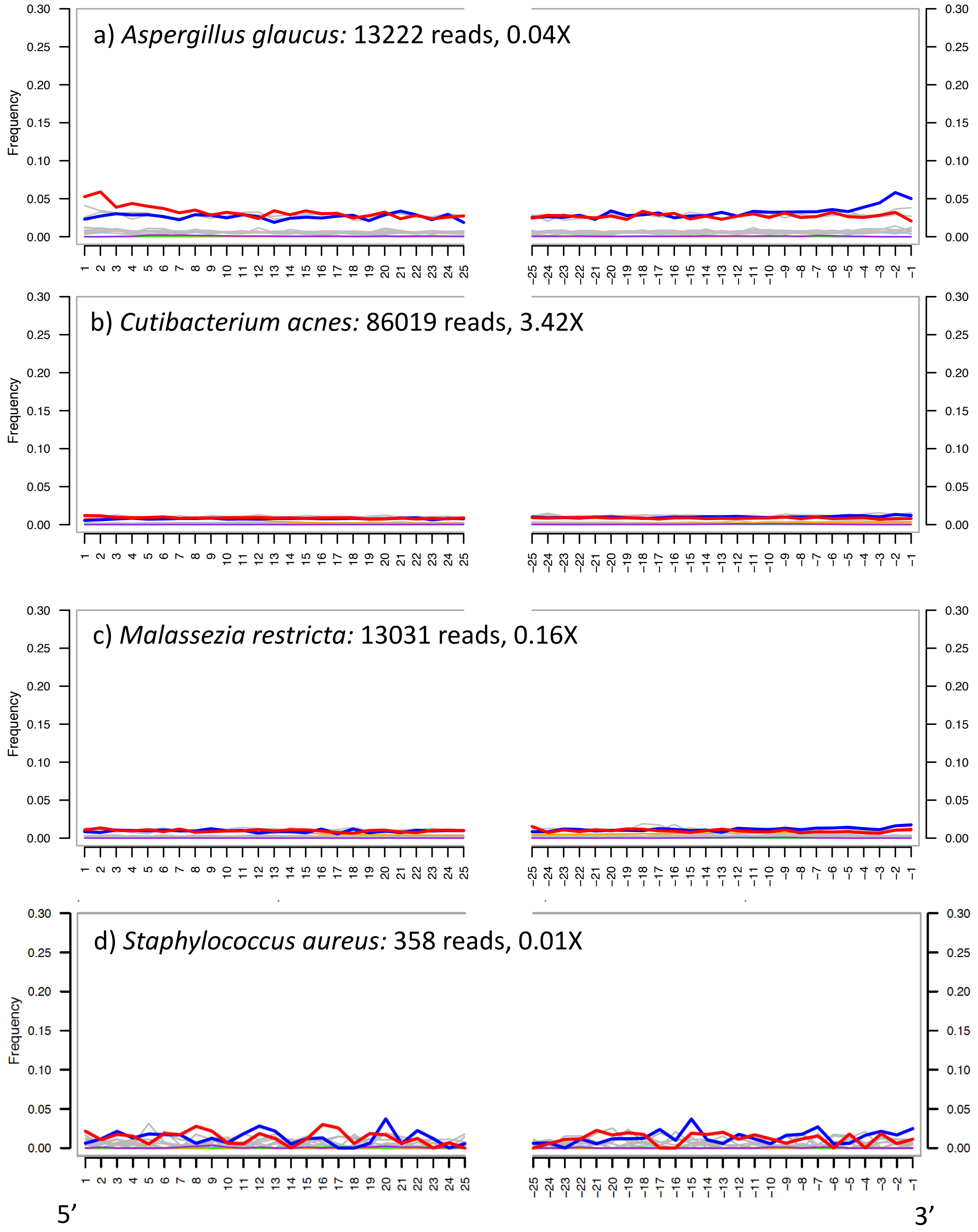


**Supplementary Figure S7:** From top to bottom; nucleotide misincorporation patterns at the ends of the DNA reads for *Aspergillus glaucus*, *Cutibacterium acnes*, *Malassezia restricta* and *Staphylococcus aureus*. The number of reads and coverage using the BWA ancient mapping pipeline is provided (see **Table S6**). Despite the low numbers of reads in some cases, the existence of this *post-mortem* deamination pattern suggests that these microbes are old. As in Supplementary Figure S2, the red line provides the C to T substitution frequency and the blue line provides the G to A substitution frequency from 5’ (left) to 3’ (right).

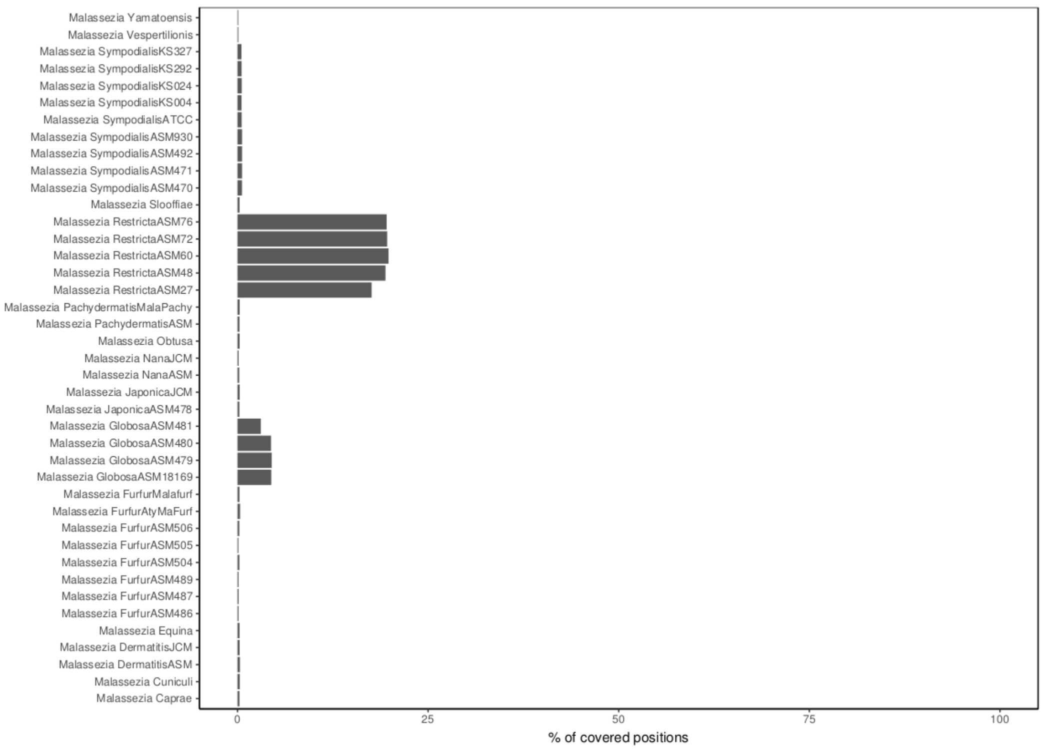


**Supplementary Figure S8:** Comparison of the percentage of genomic positions covered (x-axis) across different species and strains of the genera *Malassezia* following mapping independently to each species. Reads from the blood stain were mapped against a set of *Malassezia spp*. assemblies (y-axis).

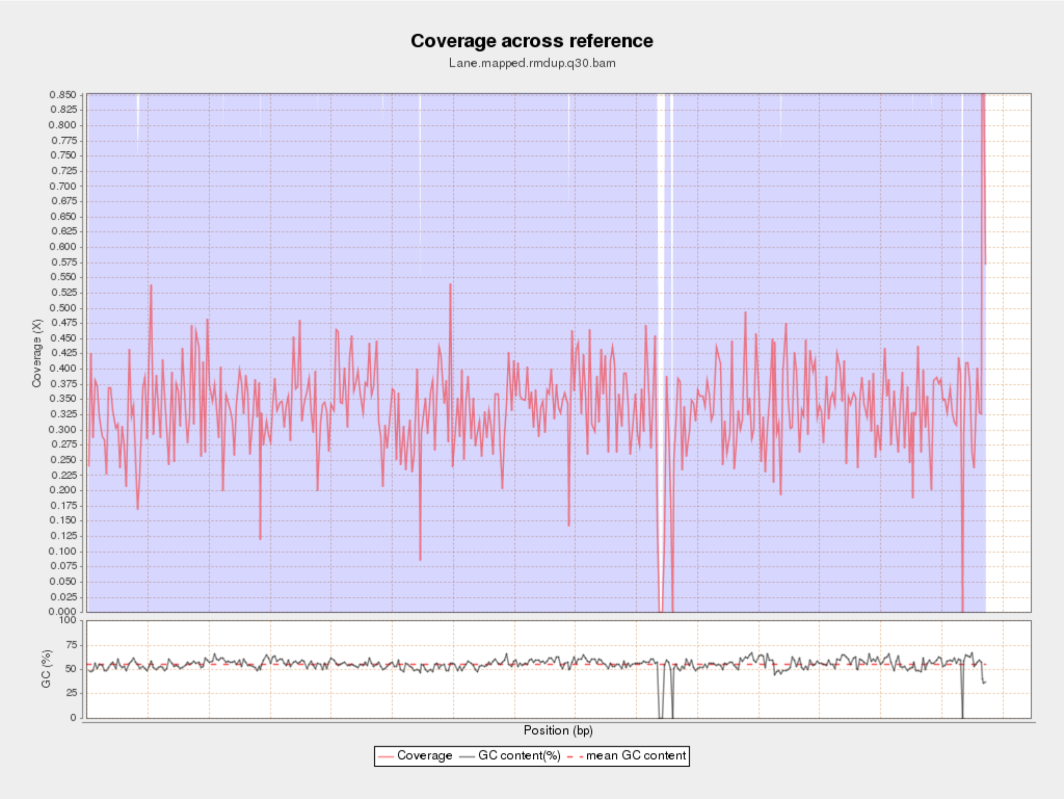


**Supplementary Figure S9:** Average coverage and GC content of reads mapped against the full (nuclear + mtDNA) *Malassezia restricta* ASM 48 reference assembly GCF_003290485.1.

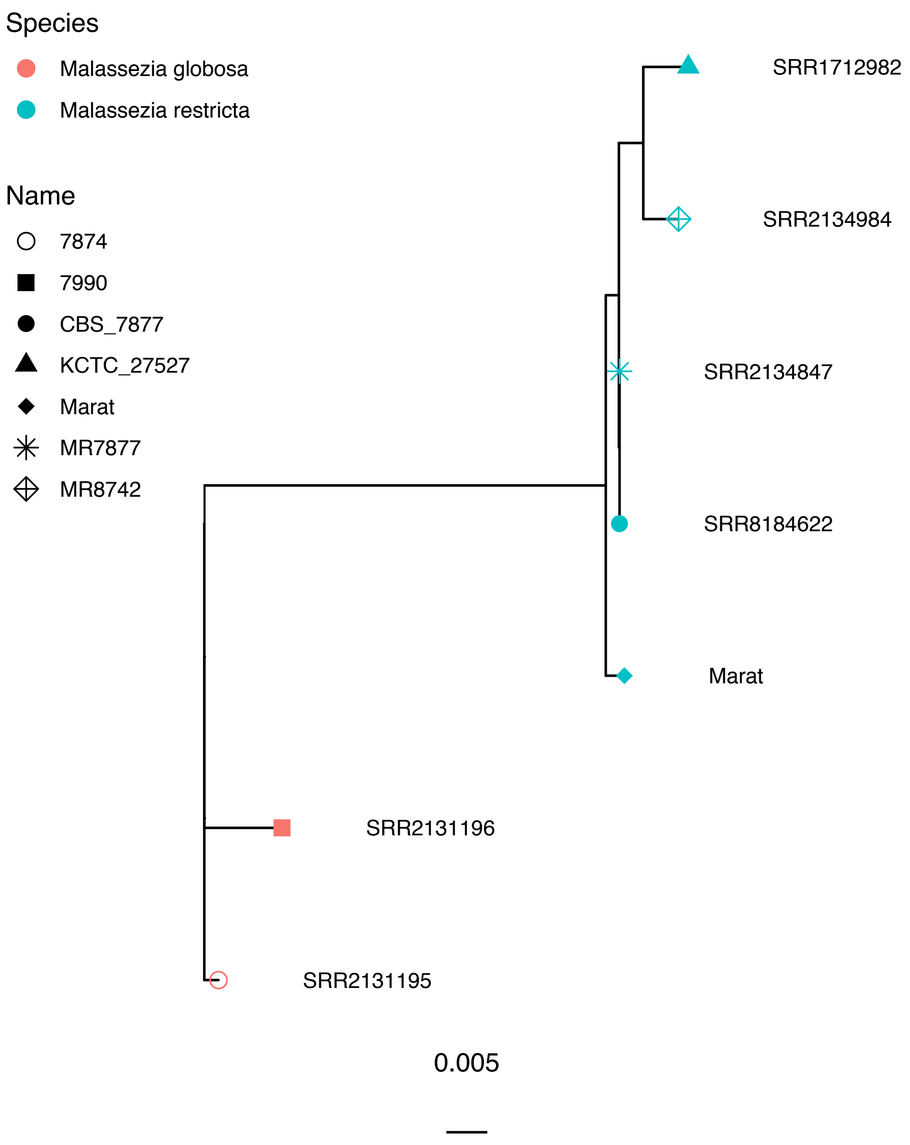


**Supplementary Figure S10:** Maximum likelihood (ML) phylogenetic tree of the *Malassezia restricta* mtDNA genome retrieved from Marat's blood stain (Marat) and four modern strains (accessions). The tree is rooted with two strains from the related *M. globosa* species.

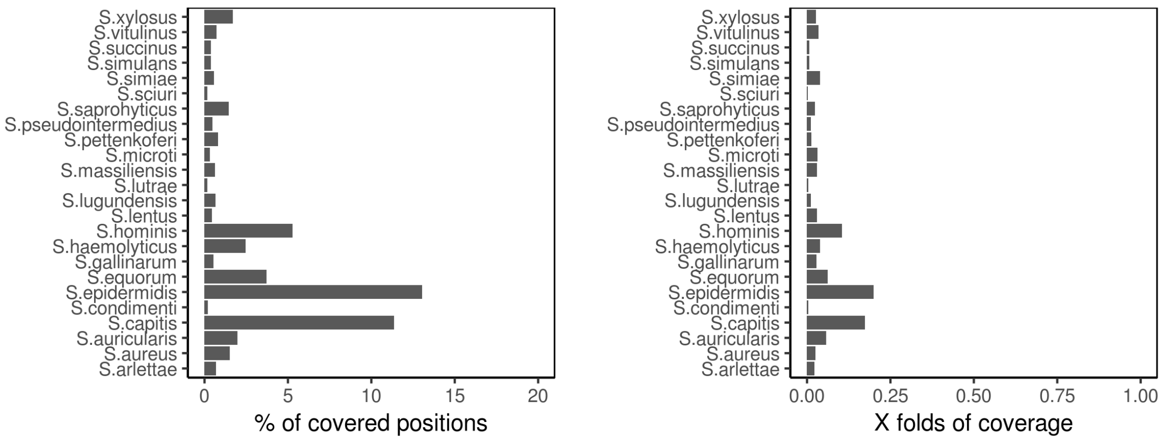


**Supplementary Figure S11:** Comparison of the percentage of genomic positions covered (x-axis) across different species and strains of the genera *Staphylococcus*. Reads from the blood stain were mapped against a set of *Staphylococcus spp*. assemblies (y-axis).

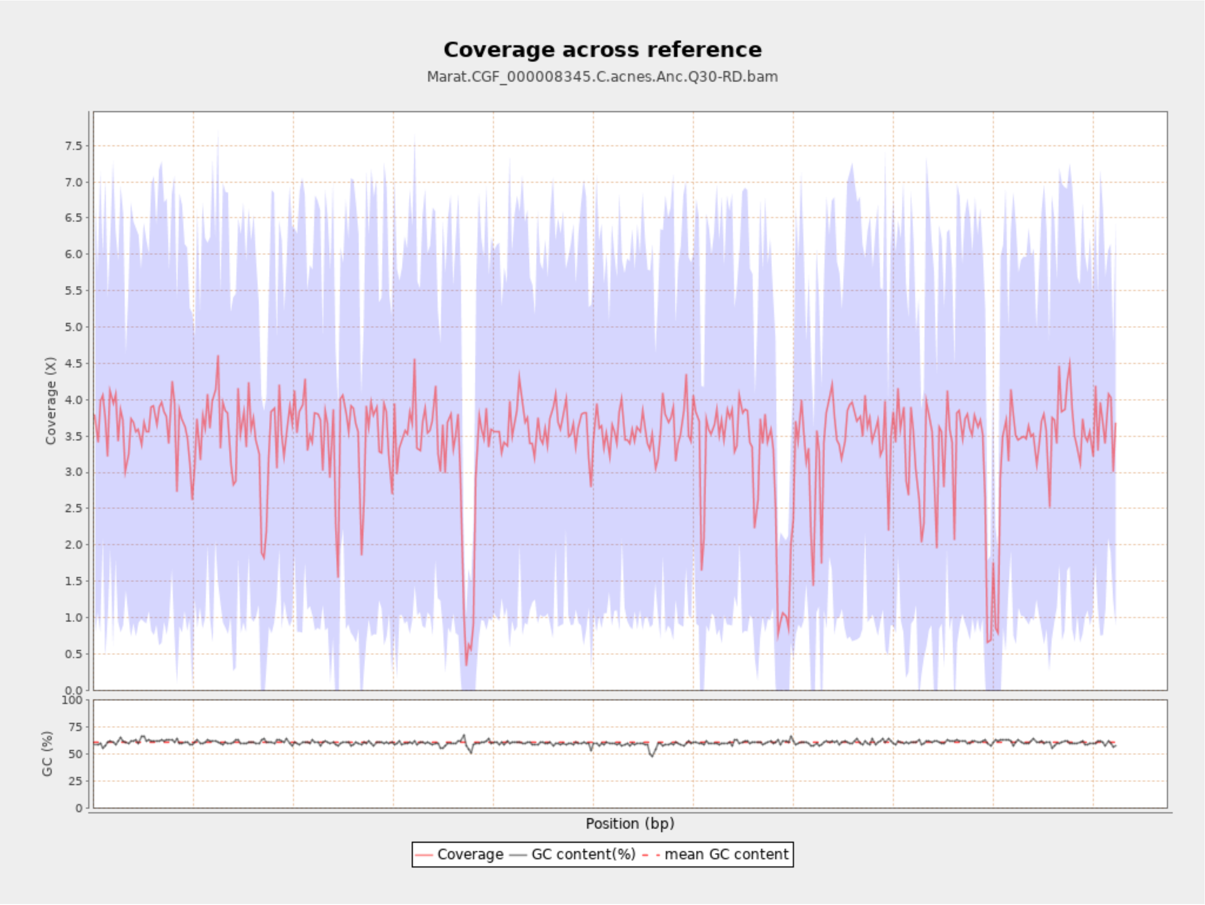


**Supplementary Figure S12:** Average coverage and GC content of all reads mapped against the *Cutibacterium acnes* ASM834v1 reference assembly.

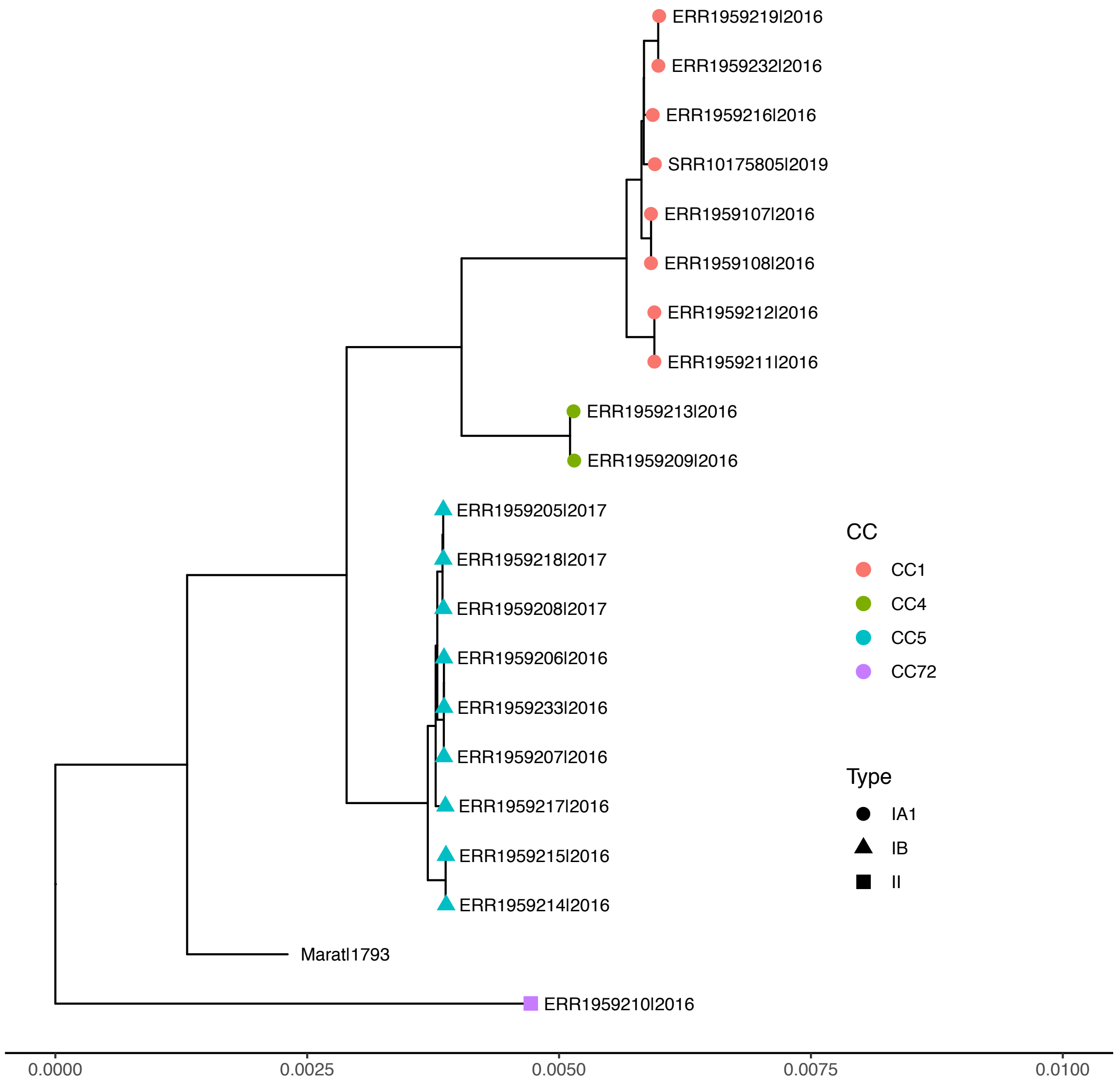


**Supplementary Figure S13:** Maximum likelihood (ML) phylogeny of the *Cutibacterium acnes* nuclear genome retrieved from Marat blood's stain together with all publicly available sequenced modern strains with median genome coverage >10X. The tree is rooted with *C. namnetense* as an outgroup (SRR9222443), with the outgroup branch not shown in the figure.
